## Supplementary material for "Anatomical basis and physiological role of cerebrospinal fluid transport through the murine cribriform plate"

**SUPPLEMENTAL FIGURES**

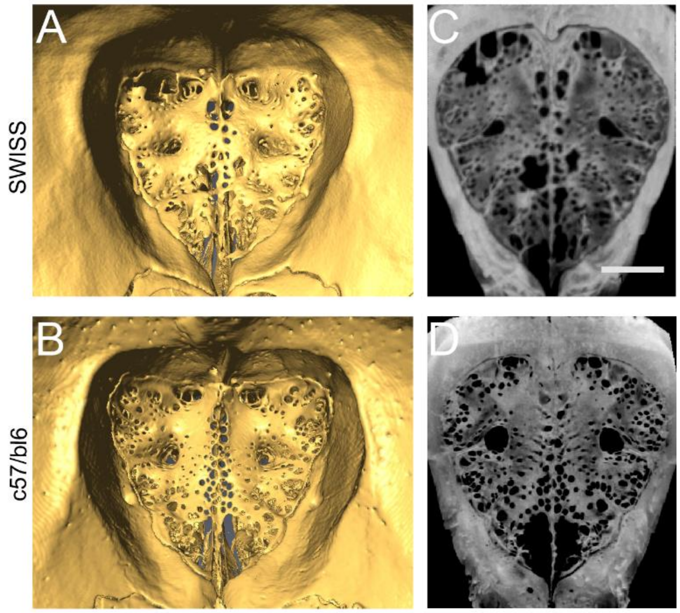


**Supplemental Figure 1**: **Cribriform plate morphology in Swiss Webster and C57BL/6J mice. A)** Reconstructed microCT image of the anterior side of the CP of a Swiss Webster mouse, showing the perforated ridge along the centerline (crista galli). Four major foramen are observed laterally from the crista galli, with one dorsal and one medial foramen on each side of the midline. **B)** Same as **(A)**, except for a C57BL/6J mouse. **C)** Max projection of the CP of a Swiss Webster mouse, note similarity to **(A)**. Scale bar 1 mm. **D)** Max projection of the CP of a C57BL/6J mouse, note similarity to **(B).** Note that the scans in **(A)** and **(B)** were taken with different scanners with different spatial resolutions: GE v|tome|x L 300 high-resolution nano/microCT scanner was used for all Swiss Webster mice, and an OMNI-X HD600 industrial microCT scanner was used for the C57BL/6J samples.

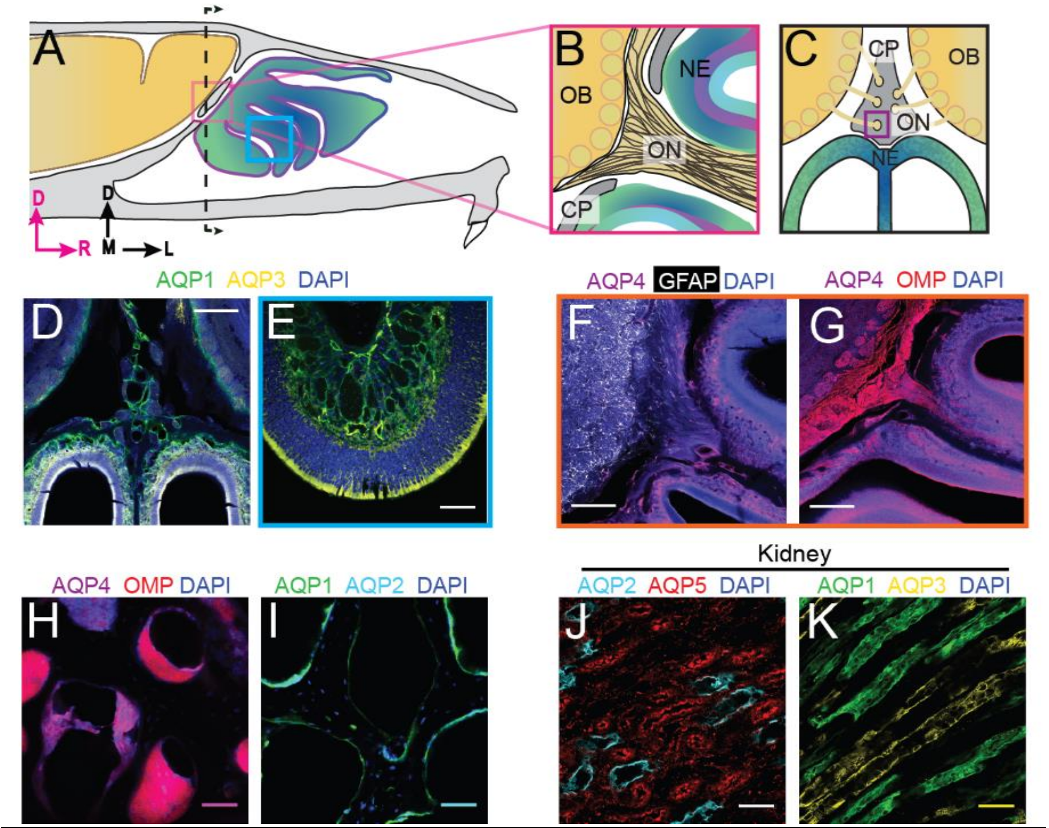


**Supplemental Figure 2: Localization of aquaporins at the olfactory nerve and bulb junction:** For schematics: olfactory nerve (ON), neuroepithelium (NE), cribriform plate (CP), olfactory bulb (OB) and glomeruli (yellow circles). D=dorsal, R=rostral, M=medial, and L=lateral. **A)** Schematic of the sagittal plane of the mouse skull and brain, showing the relationship of the OB and nerve junction to the CP. **B)** Sagittal view of the area within the pink box in **(A)**, depicting OSNs crossing the CP and terminating in the OB and glomeruli. **C)** Coronal view of the black dashed line in **(A)** illustrating the location of the CP relative to the OBs and NE. **D)** Immunofluorescent staining of area in **(C),** demonstrating the expression of AQP1, but not AQP3, along the periphery of the OB and lining the foramina of the CP. Both AQP1 and AQP3 were observed lining the lamina propria of the NE. **E)** Area depicted by the blue box in **(A),** expression of AQP1 and AQP3 lining the lamina propria of the NE. **F-G)** AQP4 (magenta), GFAP (white), OMP (red), and DAPI (blue). Examining the expression of AQP4 at the junction of the medial OB and ON and the lamina propria of the NE. **F)** Expression of AQP4 is observed lining the lamina propria of the NE and co-localizing with GFAP along the ON. **G)** Expression of AQP4 is observed lining the lamina propria of the NE and not co-localizing with OMP. **H-I)** Area indicated by purple box in **(C)**. **H)** Expression of AQP4 (magenta) is not observed lining foramina of the CP. **I)** Expression of AQP1 (green) and AQP2 (light blue) in foramina of the CP. **J-K)** Immunofluorescent staining of mouse kidney tissue. **J)** Expression of AQP2 (light blue) and AQP5 (red) in the renal cortex of the kidney. **K)** Expression of AQP1 (green) and AQP3 (yellow) in the renal medulla of the mouse kidney. **D-G)** Scale bars 250 μm. **H-K)** Scale bars 50 μm.

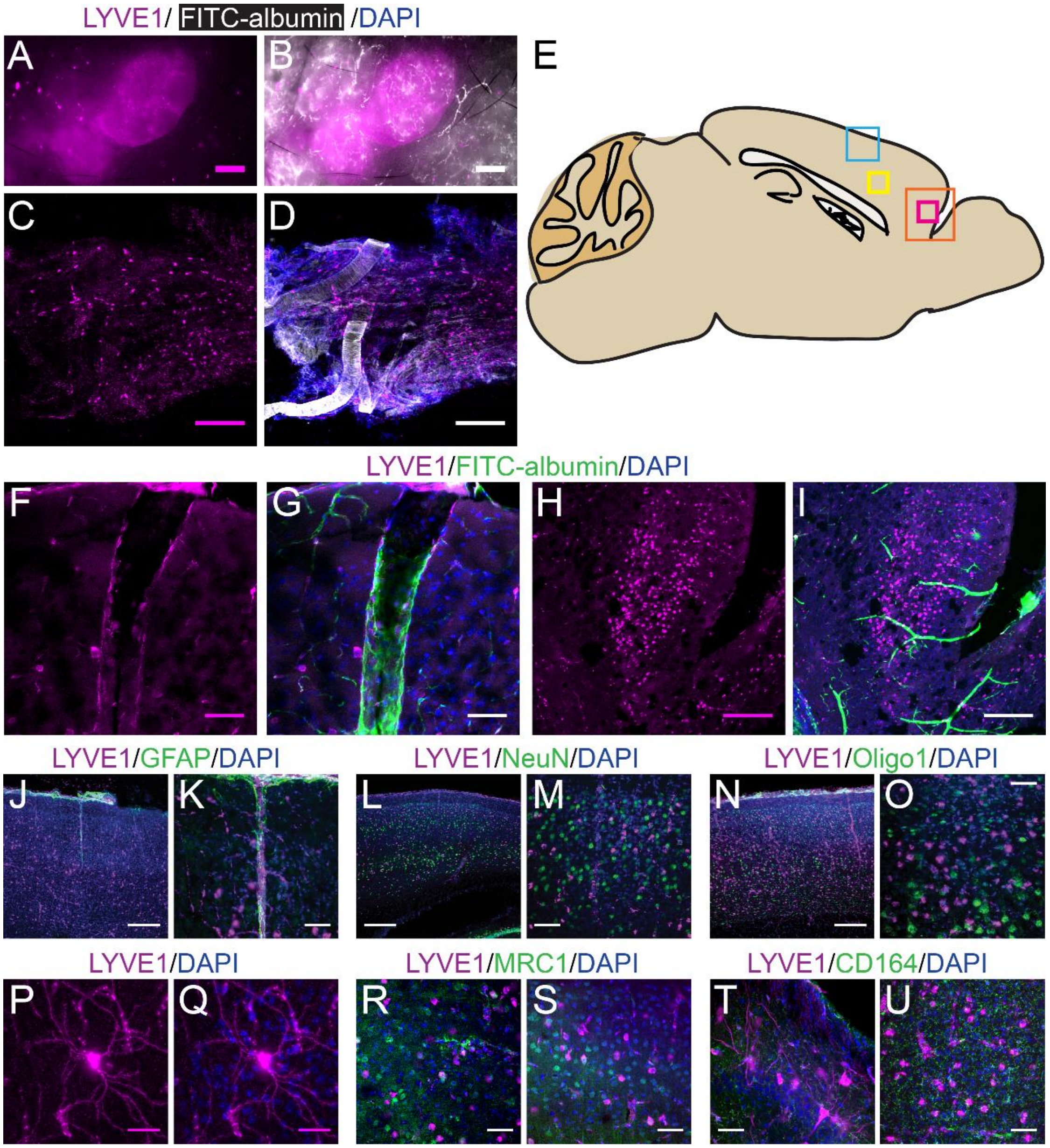


**Supplemental Figure 3: Lymphatic vessels in lymph nodes and meninges express LYVE1, as do non-lymphatic vessels and cells in the brain. A-D)** LYVE1-GFP (Ai6) mice were used for these experiments. **A-B)** Superficial lymph nodes in the neck: DiI (white) and LYVE1 (magenta). No cross-labelling is observed between $DiI^+$ blood vessels and $LYVE1^+$ lymphatic vessels. **C-D)** Meninges: DiI (white), LYVE1 (yellow), and DAPI (blue). **E)** Schematic of the sagittal view of the mouse brain showing the ROIs for (**F-O**). **F-I)** LYVE1-tdtomato (Ai14) mice were used for these experiments. FITC-albumin filled LYVE1-tdtomato mice. LYVE1 (magenta), FITC-albumin (green), and DAPI (blue). **F-G)** A penetrating vessel (FITC-albumin filled) with $LYVE1^+$ vascular endothelial cells **(F)** in the upper layers of the cortex, area indicated by the blue box in **(A)**. **H-I)** A non-vascular cell of unknown type expressing *LYVE1* in the frontal cortex **(H)**, area indicated by orange box in **(A)**. **J-U)** Immunofluorescence staining of GFAP, NeuN, Oligo1, MRC1, or CD164 in a LYVE1-GFP (Ai6) mouse in the cortex, area indicated by the blue box in **(E)**, unless specified. **J)** No overlap was observed between $GFAP^+$ and $LYVE1^+$ cells. **K)** Zoomed in area in **(J)**. **L)** No overlap was observed between NeuN$^+$ and LYVE1$^+$ cells. **M)** Zoomed in area in **(L)**. **N)** No overlap was observed between Oligo2$^+$ and LYVE1$^+$ cells. **O)** Zoomed in area in **(N)**. **P-Q)** LYVE1$^+$ cell of unknown type, area indicated by yellow box in **(E)**. **R)** Minimal overlap was observed between MRC1$^+$ and LYVE1$^+$ cells in the area indicated by the pink box in **(E).** **S)** Minimal overlap was observed between MRC1$^+$ and LYVE1$^+$ cells in the area indicated by the yellow box in **(E). T)** No overlap was observed between CD164$^+$ and LYVE1$^+$ cells in the area indicated by the pink box in **(E). U)** No overlap was observed between CD164$^+$ and LYVE1$^+$ cells in the area indicated by the yellow box in **(E). A)** Scale bar 500 μm. **C-D, H-J, L, N)** Scale bar 250 μm. **F-G, K, M, O)** Scale bar 50 μm. **P-Q)** Scale bar 25 μm.

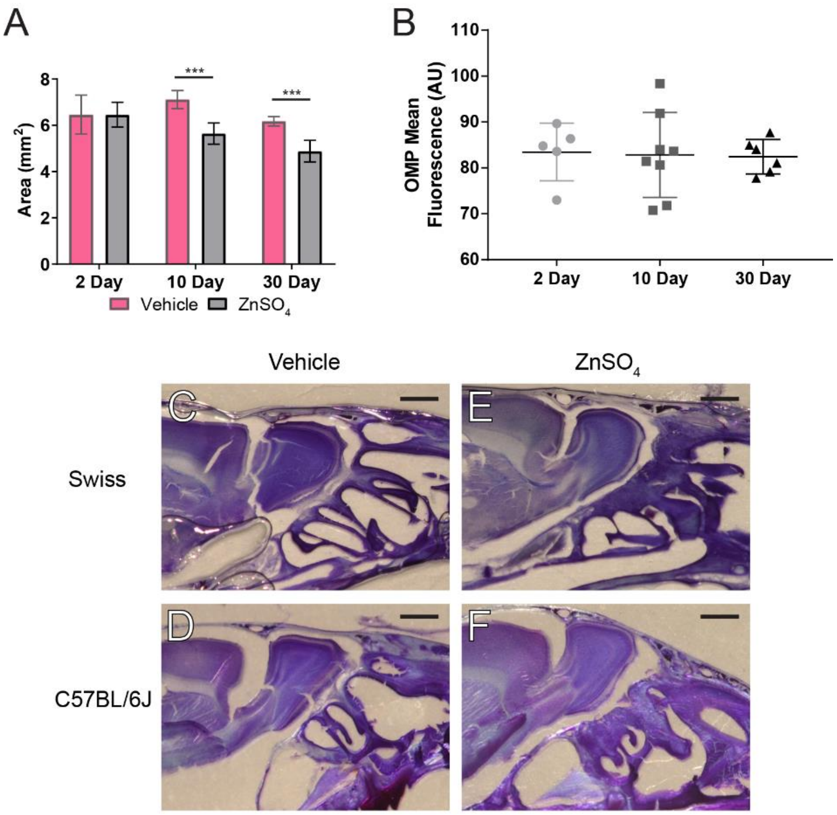


**Supplemental Figure 4: $ZnSO_4$-treatment causes olfactory bulb degeneration. A)** Comparison of the area of olfactory bulb for vehicle control and $ZnSO_4$ treated 2 days ($t(5) = 0.0073$, $p = 0.9945$, n = 3 for vehicle and n = 4 treated, post-hoc ttest2), 10 days ($t(12) = 6.46$, $p \leq 0.001$, n=7 for each group, post-hoc ttest2), and 30 days ($t(12) = 6.712$, $p \leq 0.001$, n=7 for each group, post-hoc ttest2) after treatment. Mean ± standard deviation plotted. **B)** Quantification of the OMP mean fluorescence of vehicle–treated mice. No significant differences were observed ($F(2, 16) = 0.02775$, $p = 0.9727$, 2 day n = 5, 10 day n = 8, 30 day n = 6, one-way ANOVA) showing a lack of olfactory degeneration. Mean ± standard deviation plotted. Circles, squares, and triangles represent means of individual animals. **(C-F)** Thionin-nissl staining 30 days post-$ZnSO_4$ treatment of Swiss Webster and C57BL/6J vehicle mice **(C-D)** and 30 days post $ZnSO_4$ treated mice **(E-F)**. Note increased space between OB and CP and change in morphology of the neuroepithelium in $ZnSO_4$-treated mice. Scale bar 1 mm.

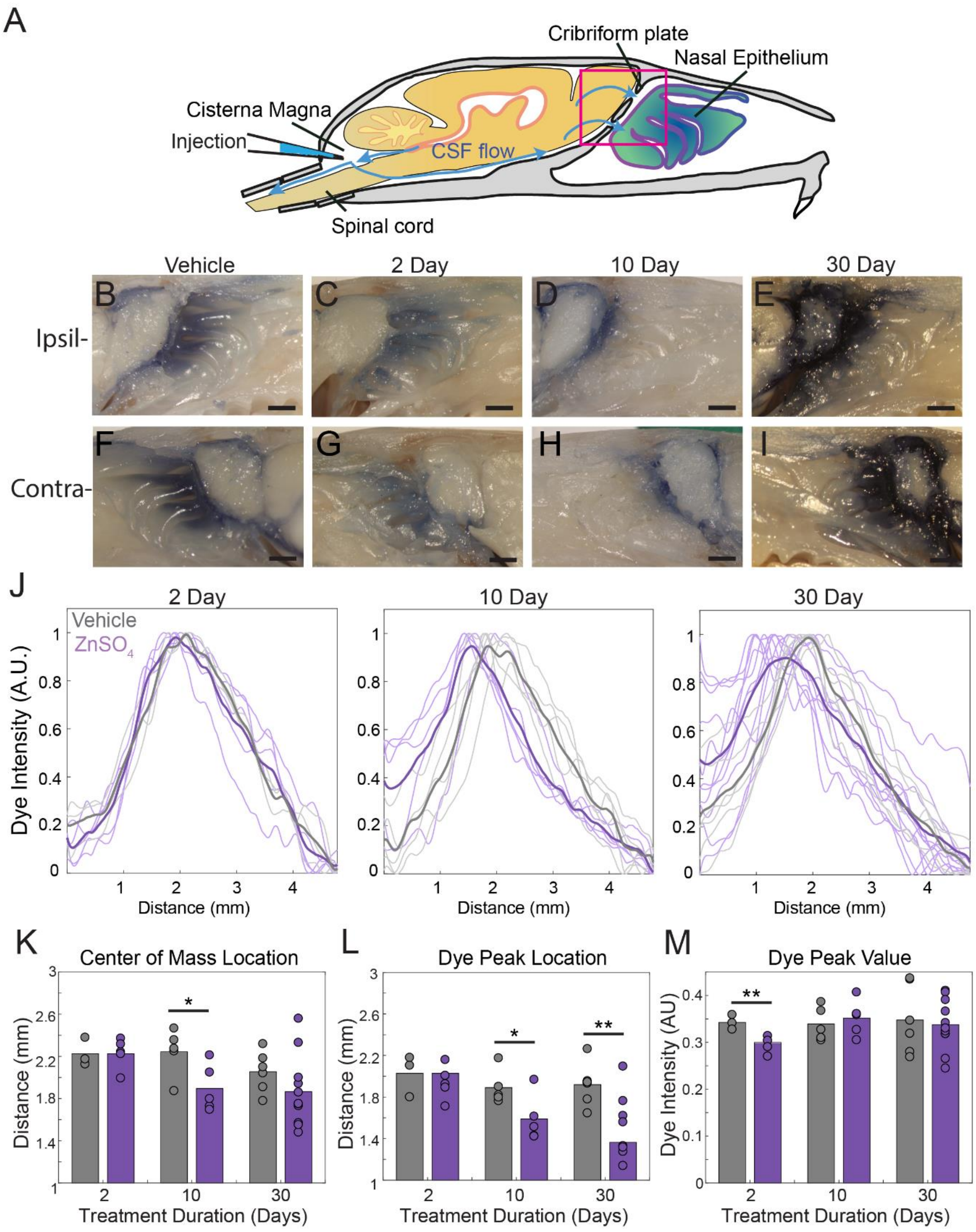


**Supplemental Figure 5: $ZnSO_4$ treatment impacts CSF outflow on the side contralateral to treatment. A**) Sagittal view of the skull depicting direction of CSF flow and location of cisterna magna injection of EB. **B-E)** Sagittal view of decalcified and cut skull of the ipsilateral side for imaging after EB injection in the cisterna magna, area indicated by black box in **(A). F-I)** Contralateral side of $ZnSO_4$ treatment. Scale bar 1mm. **B, F)** Vehicle control. **C, G)** 2 days post-$ZnSO_4$ treatment. **D, H)** 10 days post-$ZnSO_4$ treatment. **E, I)** 30 days post-

$ZnSO_4$ treatment. **J)** Comparison of all dye intensity curves of all vehicle (gray) and $ZnSO_4$ treated (purple) animals for each treatment duration: 2 day, n = 3 vehicle and n = 5 treated. 10 day, n = 5 for each group. 30 day, n= 6 vehicle and n = 11 for treated. Means of the group plotted as thicker lines. **K-M)** Mean is plotted as height of bar. Circles represent individual animals. **K)** Mean center of mass of the dye intensity on the side contralateral to the treatment for vehicle and zinc treated animals after 2 ($t(6) = -0.0290$, $p = 0.9978$, n = 3 vehicle and n = 5 treated, post-hoc ttest2), 10 ($t(8) = 2.4522$, $p = 0.0398$, n = 5 for each group, post-hoc ttest2), and 30 ($t(15) = 1.2746$, $p = 0.2218$, n= 6 vehicle and n = 11 for treated, post-hoc ttest2) days of treatment. **L)** Mean dye peak location of the contralateral side plotted for vehicle and zinc treated animals after 2 ($t(6) = 0.6881$, $p=0.5171$, n = 3 vehicle and n = 5, post-hoc ttest2), 10 ($t(8) = 2.4056$, $p = 0.0428$, n = 5 for each group, post-hoc ttest2), and 30 ($t(15) = 3.4097$, $p = 0.0039$, n= 6 vehicle and n = 11 for treated, post-hoc ttest2) days of treatment. **M)** Maximum dye peak value of the contralateral side plotted for vehicle and $ZnSO_4$ treated animals after 2 ($t(6) = -3.8517$, $p = 0.0084$, n = 3 vehicle and n = 5 treated, post-hoc ttest2), 10 ($t(8) = 0.5298$, $p = 0.6107$, n = 5 for each group, post-hoc ttest2), and 30 ($t(15) = -0.3148$, $p = 0.7573$, n = 6 vehicle and n = 11 for treated, post-hoc ttest2) days of treatment. *$p\leq0.05$. **$p\leq0.01$.

__Supplementary Tables__

| Evans blue drainage | | | | | |
|---|---|---|---|---|---|
| **F** | **d.o.f.** | **p** | **Side** | **Parameters** | **Parameters** |
| *5.45* | *2* | ***0.0086*** | *IPSI* | *CoM* | *Interaction: duration * type* |
| 0.49 | 2 | 0.6144 | IPSI | PL | Interaction: duration * type |
| 0.61 | 2 | 0.5494 | IPSI | PV | Interaction: duration * type |
| *11.05* | *2* | ***0.0002*** | *IPSI* | *CoM* | *effect: duration* |
| *13.16* | *1* | ***0.0009*** | *IPSI* | *CoM* | *effect: type* |
| *3.63* | *2* | ***0.0365*** | *IPSI* | *PL* | *effect: duration* |
| *13.72* | *1* | ***0.0007*** | *IPSI* | *PL* | *effect: type* |
| *5.27* | *2* | ***0.0098*** | *IPSI* | *PV* | *effect: duration* |
| 0 | 1 | 0.9497 | IPSI | PV | effect: type |
| 1.06 | 2 | 0.3601 | CONTRA | CoM | Interaction: duration * type |
| 2.08 | 2 | 0.1435 | CONTRA | PL | Interaction: duration * type |
| 0.79 | 2 | 0.4645 | CONTRA | PV | Interaction: duration * type |
| 2.93 | 2 | 0.0694 | CONTRA | CoM | effect: duration |
| 3.84 | 1 | 0.0596 | CONTRA | CoM | effect: type |
| *4.19* | *2* | ***0.0253*** | *CONTRA* | *PL* | *effect: duration* |
| *10.64* | *1* | ***0.0028*** | *CONTRA* | *PL* | *effect: type* |
| 0.73 | 2 | 0.49 | CONTRA | PV | effect: duration |
| 0.67 | 1 | 0.4188 | CONTRA | PV | effect: type |

**Table 1:** Two-way ANOVA statistics for all Evans blue cisterna magna injections. d.o.f. = degrees of freedom. CoM = center of mass, PL = peak location, PV = peak value. All significant p values are bolded ($p \leq 0.05$).

| Intranasal and kaolin treatment effects on intracranial pressure (ICP) | | | | | | | | |
|---|---|---|---|---|---|---|---|---|
| Treatment | ICP mmHg (vehicle) | ICP mmHg (treated) | t | d.o.f. | p | n (vehicle) | n (treated) | assay |
| Vehicle/$ZnSO_4$ | 3.40 ± 1.24 | 5.34 ± 1.09 | 2.1408 | 5 | 0.0852 | 4 | 3 | ICP Rest. 60 day |
| Vehicle/$ZnSO_4$ | 7.28 ± 2.69 | 7.95 ± 1.91 | 0.3606 | 5 | 0.7331 | 4 | 3 | ICP Locomotion. 60 day |
| *aCSF/kaolin* | *9.14 ± 2.12* | *21.54 ± 2.98* | *5.88* | *4* | ***0.0042*** | *3* | *3* | *ICP Locomotion* |
| aCSF/kaolin | 4.10 ± 2.26 | 11.35 ± 6.50 | 1.82 | 4 | 0.1423 | 3 | 3 | ICP Rest. |

**Table 2:** Unpaired t-test results for intracranial pressure (ICP) measurements. d.o.f. = degrees of freedom. All significant p values are bolded ($p \leq 0.05$).

| Intranasal treatment effects on intracranial pressure (ICP) | | | | | |
|---|---|---|---|---|---|
| ICP mmHg (vehicle) | ICP mmHg(treated) | p | n (vehicle) | n (treated) | assay |
| 5.72 ± 4.27 | 4.12 ± 2.04 | 0.5949 | 11 | 9 | ICP Rest. 10 day |
| 5.94 ± 4.31 | 3.13 ± 1.20 | 0.3429 | 4 | 4 | ICP Rest. 30 day |
| 8.77 ± 5.17 | 6.36 ± 2.10 | 0.5949 | 11 | 9 | ICP Locomotion. 10 day |
| 9.21 ± 5.04 | 6.23 ± 1.39 | 0.6857 | 4 | 4 | ICP Locomotion. 30 day |

**Table 3**: Mann-Whitney U test statistics for intracranial pressure (ICP) measurements.
